# Adenovirus transduction to express human ACE2 causes obesity-specific morbidity in mice, impeding studies on the effect of host nutritional status on SARS-CoV-2 pathogenesis

**DOI:** 10.1101/2021.05.26.445786

**Authors:** Pallavi Rai, Christina Chuong, Tanya LeRoith, James W Smyth, Julia Panov, Moshe Levi, Kylene Kehn-Hall, Nisha K. Duggal, James-Weger Lucarelli

## Abstract

The COVID-19 pandemic has paralyzed the global economy and resulted in millions of deaths globally. People with co-morbidities like obesity, diabetes and hypertension are at an increased risk for severe COVID-19 illness. This is of overwhelming concern because 42% of Americans are obese, 30% are pre-diabetic and 9.4% have clinical diabetes. Here, we investigated the effect of obesity on disease severity following SARS-CoV-2 infection using a well-established mouse model of diet-induced obesity. Diet-induced obese and lean control C57BL/6N mice, transduced for ACE2 expression using replication-defective adenovirus, were infected with SARS-CoV-2, and monitored for lung pathology, viral titers, and cytokine expression. No significant differences in tissue pathology, viral replication or cytokine expression were observed between lean and obese groups. Notably, significant weight loss was observed in obese mice treated with the adenovirus vector, independent of SARS-CoV-2 infection, suggesting an obesity-dependent morbidity induced by the vector. These data indicate that the adenovirus-transduced mouse model of SARS-CoV-2 infection is inadequate for performing nutrition studies, and caution should be used when interpreting resulting data.

## Introduction

Coronavirus disease-2019 (COVID-19) is the third pandemic in the 21^st^ century caused by a novel coronavirus, after severe acute respiratory syndrome coronavirus (SARS-CoV) in 2003 [1,2,3,4] and the Middle East respiratory syndrome coronavirus (MERS-CoV) in 2012 [5,6,7]. COVID-19 is responsible for ~99 million confirmed cases, with over 2 million deaths globally, and ~25 million confirmed cases and ~400,000 deaths in the U.S. alone, as of January 26^th^, 2021 [8]. COVID-19 is characterized by fever and respiratory symptoms, which can progress to more severe and fatal disease [9]. Infection with SARS-CoV-2, the causative agent of COVID-19, is more likely to cause critical illness or deaths in the elderly, immunocompromised and individuals with co-morbidities such as obesity, diabetes, and hypertension [10,11,12,13].

Obesity and diabetes have been identified as risk factors for poor outcomes in other major pandemics involving respiratory tract infections like MERS [14] and influenza [15,16,17]. Like COVID-19, obesity too has become pandemic, affecting 12.5% of people globally and approximately 42% of Americans [18]. Obesity is the most frequently observed component of metabolic syndrome, a condition that clusters abdominal obesity, dyslipidemia (abnormally high blood lipid levels), hyperglycemia (increased blood sugar levels) and hypertension (increased blood pressure) [19]. Experiments using diet-induced obese mice to study the pathogenesis of influenza A virus have shown not only are obese mice more susceptible to infection [20] but that they also promote the emergence of more virulent strains of the virus [21]. To study the impact of obesity on SARS-CoV-2 pathogenesis and evolution, a similar mouse model is required; however, mice are not susceptible to infection with SARS-CoV-2 due to the lack of human Angiotensin Converting Enzyme 2 (hACE2) receptor, required for entry of the virus [22,23]. Several strategies have been developed to circumvent this issue: transgenic mice expressing hACE2 (hACE2-Tg) [24,25,26,27], mouse-adapted SARS-CoV-2 strains [28,29], and transduction with adenovirus vectors expressing hACE2 (AdV-hACE2) [30,31]. The AdV-hACE2 approach has several advantages: it can be used with different strains of mice, is easy to manipulate, causes little pathology to wild-type mice, and allows for the use of wild-type (WT) SARS-CoV-2 [30,31].

In this study, we sought to investigate the effect of obesity on disease outcome following SARS-CoV-2 infection using an adenovirus-transduced mouse model. Currently, no mouse models for studying the impact of nutritional status on COVID-19 have been described, limiting progress towards understanding the mechanisms leading to more severe disease in people with co-morbidities. To this end, diet-induced obese and lean mice were transduced with AdV-hACE2 followed by SARS-CoV-2 infection and monitored for morbidity, mortality, virus replication and cytokine expression. Surprisingly, obese, but not lean, mice inoculated with the replication-defective AdV-hACE2 lost significant weight, independent of SARS-CoV-2 infection, suggesting an obesity-dependent morbidity induced by the AdV-hACE2 vector. SARS-CoV-2 replication occurred in the lungs of both lean and obese mice, though, we did not observe differences in terms of disease outcomes, viral replication, or cytokine expression when normalized to mock-SARS-CoV-2-infected controls. Accordingly, due to the confounding influence of AdV-hACE2 on SARS-CoV-2 infection, we could not determine obesity’s impact on SARS-CoV-2 infection. These data underscore the importance of selecting an appropriate model for nutrition studies and suggest that researchers should exercise caution when using the AdV-hACE2 model to study SARS-CoV-2. Further studies using transgenic mice, mouse-adapted strains [28,29], or other animals such as hamsters [32,33] or ferrets [34,35] should be conducted to explore the relationship between COVID-19 and metabolic co-morbidities.

## Materials and Methods

### Cells and Viruses

Vero E6 cells were obtained from American Type Culture Collection (ATCC; CRL-1586) and were maintained in Dulbecco’s Modified Eagle’s Medium (DMEM) supplemented with 5% fetal bovine serum (FBS), 1mg/mL gentamicin, 1% non-essential amino acids (NEAA) and 25 mM HEPES buffer in an incubator at 37°C with 5% CO_2_.

The replication-defective adenovirus encoding human ACE2 (AdV-hACE2) was originally prepared by the University of Iowa and Washington University in St. Louis, as reported previously [3]. The viral stocks were propagated using 293A cells (ThermoFisher Catalog # R70507) and purified by cesium chloride gradient centrifugation prior to titering, also in 293A cells, as previously described [36]. SARS-CoV-2 strain 2019 n-CoV/USA_WA1/2020 was obtained from BEI Resources (Catalog # NR-52281) [37] and was propagated in Vero E6 cells to prepare stocks; infectious titers were determined by plaque assays on the same cell line.

### Mice and Diets

C57BL/6N male and female mice were obtained from Charles River Laboratories at three to four weeks of age and allowed to acclimatize for a week before initiating diets. Mice were housed in groups of five per cage and maintained at ambient temperature with *ad libitum* supply of food and water, except for overnight food deprivation (12-14 hours) prior to blood glucose measurements. All animal handling protocols were approved by the Institutional Animal Care and Use Committee (Protocol #20-060) at Virginia Tech.

All diets used for the study were obtained from Research Diets (New Brunswick, NJ, USA). Twenty mice (10 males and 10 females) were placed on a low-fat diet with 10% kcal fat (LFD; D12450K) and 20 (10 males and 10 females) on a high-fat diet with 60% kcal fat (HFD; D12492). Throughout the manuscript, we will refer to the groups as follows: lean (low fat diet) or obese (high fat diet). The mice were kept on these diets for 18-20 weeks before infections, and the same diets were continued until the end of the experiment. Table S1 displays the caloric information of these diets.

### Fasting glucose levels

Two weeks prior to transduction by the adenovirus (16 weeks after diet initiation), mice were deprived of food overnight (12-14 hours), blood was collected by submandibular bleed and glucose levels were measured by Abox glucose monitoring kit.

### Mouse infections

Mice were moved to a BSL-3 facility 24 hours prior to SARS-CoV-2 infections. For primary infections with the AdV-hACE2, 19-20-week-old mice were anesthetized with ketamine and xylazine (90 mg/kg and 5 mg/kg respectively) intraperitoneally and then inoculated intranasally with 10^8^ plaque-forming units (PFUs) of the virus in 50 μL of Roswell Park Memorial Institute (RPMI)-1640 media with no additives. Five days post-adenovirus inoculation, the mice received 10^5^ PFUs of SARS-CoV-2 intranasally under ketamine and xylazine anesthesia, or RPMI-1640 media alone for mock-infected groups. Six infected and four control mice from each group were sacrificed at days 4 and 10 after SARS-CoV-2 infection. Mice were weighed daily post-SARS-CoV-2 infection and monitored for visual symptoms of the disease. Blood was collected via submandibular bleeds (~200μL per mouse), serum was collected via centrifugation at 5000 x g for 10 minutes, transferred to fresh tubes and stored at −80°C.

### Histopathology and organ titration

Tissues for viral titration were harvested aseptically in 2 mL tubes containing a 5 mm stainless steel bead, RPMI-1640, 10 mM HEPES and 1% FBS (herein referred to as the “tissue diluent”), to a final concentration of 10% weight by volume. (QIAGEN) at 30 cycles per second for 2 mins and then centrifuged at 5000 x g for 10 mins. Plaque assays were performed on the clarified supernatant. Briefly, Vero E6 cells at 100% confluency were inoculated with 50 μL of the serially diluted samples and incubated at 37°C in 5% CO_2_ for 1 hour. Overlay media containing 0.6% tragacanth gum, 1x MEM (minimum essential media), 20 mM HEPES and 4% FBS were then added, and the plates were allowed to incubate for 2 days for plaque formation.

For histopathology, the tissues harvested from mice were fixed in 4% paraformaldehyde for at least 1 week. The Virginia Tech Animal Laboratory Services (ViTALS) performed paraffin embedding, sectioning and hematoxylin-eosin staining, and a board-certified pathologist scored the slides in a blinded manner.

### RNA extraction

A section of the infected lung tissues (4 dpi) was collected in 0.5 mL of TRIzol LS reagent (ThermoFisher) in 2 mL tubes with a 5mm bead. Lung tissues were homogenized using a TissueLyser II (QIAGEN) at 30 cycles per second for 2 mins and stored at −80°C. Samples were thawed and total RNA was extracted using the manufacturer’s protocol for TRIzol extraction.

### Reverse Transcription Quantitative PCR (RT-qPCR)

To quantify SARS-CoV-2 genomes, the 2019-nCoV RUO kit from Integrated DNA Technologies (IDT, Leuven, Belgium) was utilized. N2 combined primer-probe mix from the kit was used (Table S2) with Quantabio qScript XLT-One-Step RT-qPCR ToughMix (2X). The Bio-Rad CFX-96 (Hercules, CA, USA) was used for RT-qPCRs with the following conditions: 50°C for 10 mins for reverse transcription, 95°C for 3 mins for initial denaturation and polymerase activation, followed by 45 cycles of 95°C for 10 secs for denaturation, and 60°C for 30 secs for annealing/extension. For generating the standard curve, we used N gene RNA generated by *in vitro transcription* and used ten-fold serial dilutions to the point where no genome was detectable by qPCR. The virus concentration was calculated by fitting the Cq values of the samples to the standard curve and expressed in terms of N-gene copies/mL.

RT-qPCR was also performed on RNA extracted from the 4 dpi lung samples with NEB Luna Universal One-Step RT-qPCR Kit with SYBR-Green (NEB, Ipswich, MA, USA) to quantify the cytokines IL-6 and IFN-β. Primers were obtained from IDT and are listed in Table S2. The conditions for the reactions in a Bio-Rad CFX-96 were: 50°C for 10 mins for reverse transcription, 95°C for 1 min for initial denaturation and polymerase activation, followed by 45 cycles of 95°C for 10 secs for denaturation, and 60°C for 30 secs for annealing/extension, followed by a melt curve. The samples were calibrated with mouse GAPDH as the reference gene with respect to mock-infected groups as control. The relative expression/fold change was calculated using the Delta-Delta-Ct (∆∆Ct) method of relative quantification [38].

### Statistical Analysis

Statistical analysis was done using GraphPad Prism 9 (GraphPad Software, San Diego, CA, USA). Details of the statistical technique used for analyses are provided in the figure legends. The level of significance has been determined by the following p-values: p=0.1234 (ns); p = 0.0332 (*), p = 0.0021 (**), p = 0.0002 (***), p<0.0001 (****). The error bars represent standard deviation (SD) from the mean, in all the figures and the dotted lines denote the limit of detection (L.O.D.). All statistical analyses were performed on data that passed the normality test using Shapiro-Wilk test.

## Results and Discussion

### Wild type C57BL/6N mice fed a high fat diet had increased weight gain and blood glucose levels

The proportion of COVID-19 patients requiring intensive care was previously reported to be directly proportional to the body-mass index (BMI), being highest in patients with BMI ≥35 (morbidly obese) [39]. Obesity has also been correlated with severe outcomes during the H1N1 influenza-A epidemic [40,41,42,17]. Furthermore, diet-induced obese mice had significantly higher mortality rates when infected with influenza virus [20] and more severe disease following chikungunya virus (CHIKV) [43] and dengue virus (DENV) infection [44]. We sought to use a similar diet-induced obese mouse model to investigate whether obese mice infected with SARS-CoV-2 exhibit worse disease outcomes as compared to their lean counterparts.

We observed a significant weight gain in high-fat diet (HFD)-fed mice compared to the low-fat diet (LFD)-fed groups starting 7 weeks after the initiation of diets (p-value <0.0001) (Fig. 1A). Weights were analyzed on the combined data for males and females on LFD and HFD. Separate analysis for males and females is presented in supplementary figure S1. To assess other metabolic parameters, we measured overnight fasting blood glucose levels after 9 and 16 weeks of the initiation of diets. We tested only 10 mice per group at 9 weeks to reduce stress on the mice. Hyperglycemia, defined as overnight fasting glucose levels of ≥15 0mg/dL, was not significantly different between the lean and obese groups at 9 weeks (p-value=0.2344) (Fig. 1B). By 16 weeks, however, there was a significant difference between HFD and LFD groups (p-value=0.0001) (Fig. 1C). This 15 to 20-week diet regimen produced a mouse model that mirrored obesity and hyperglycemia in humans and, therefore, was appropriate for infection studies.

**Fig.1:**
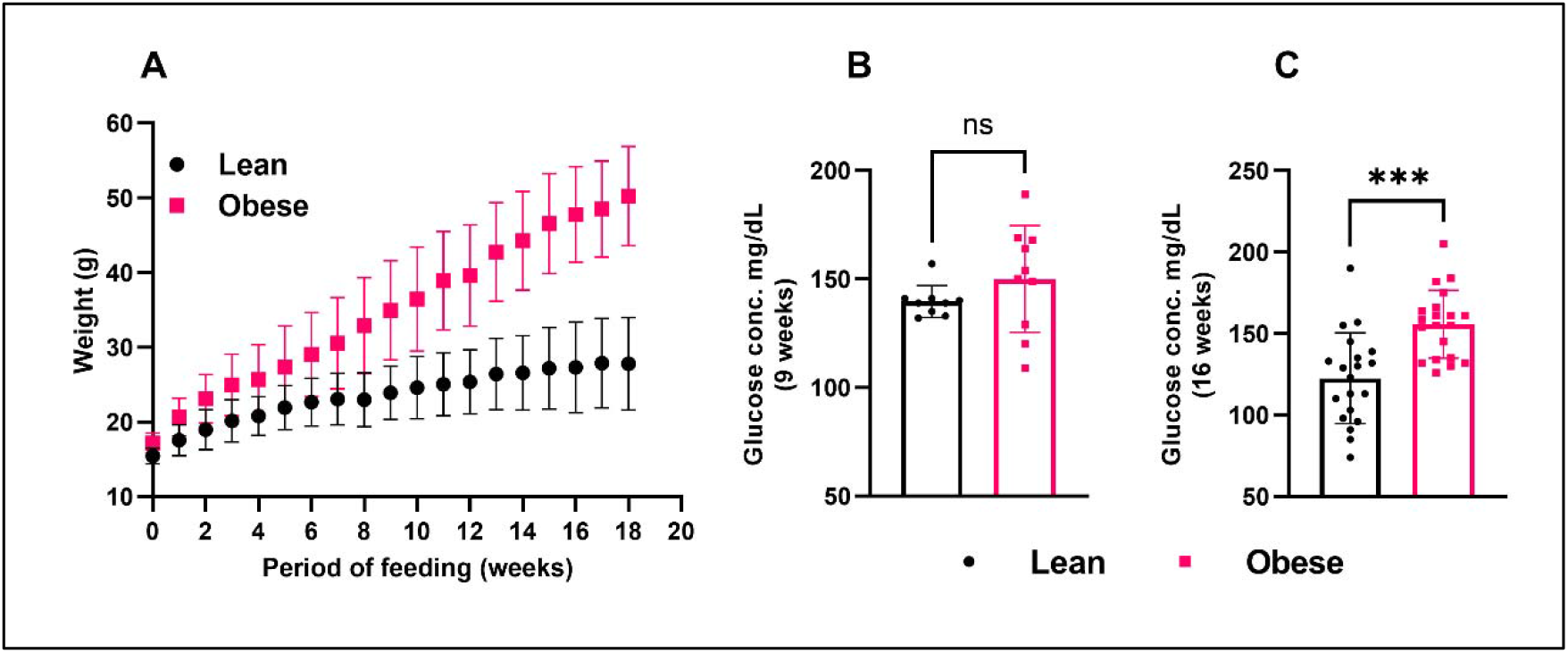
Metabolic characterization of mice fed low fat (lean) and high fat diets (obese). **A)** Four-week-old C57BL/6N mice (20/group) were fed a high-fat diet (60% fat) or a low-fat diet (10% fat) for 15-20 weeks. Weights were measured weekly after initiating the diets. Statistical analysis was done using 2way-ANOVA with Sidak’s multiple comparisons test. **B)** Blood glucose levels were measured following nine weeks of feeding **C)** Glucose levels were measured after 16 weeks of feeding. Statistical analysis was done using Mann-Whitney test and a significantly high level of glucose was observed in obese mice compared to lean ones after 16 weeks of diet-initiation (p-value=0.0001). The error bars indicate standard deviation (SD) from mean.

### Adenovirus-transduction caused morbidity in obese mice, independent of SARS-CoV-2 infection

Since mouse ACE2 receptors do not allow for WT SARS-CoV-2 infection, we first transduced mice with a replication-defective adenovirus encoding for human ACE2 (AdV-hACE2) intranasally to achieve hACE2 expression in the lungs and render mice susceptible to infection. We then inoculated the mice with SARS-CoV-2 or diluent five days post-transduction, as previously described [30,31] (Fig. 2A). Surprisingly, the obese mice inoculated only with AdV-hACE2 showed significant weight loss, similar to those infected with both AdV-hACE2 and SARS-CoV-2 (Fig. 2B). No weight loss was observed in lean mice treated with AdV-hACE2 or AdV-hACE2 and SARS-CoV-2. We found a significant weight loss in obese-AdV only mice compared to the lean-AdV only group starting from days 5 (p-value=0.0010) through day 9 post-AdV transduction (p-value=0.0242). The weights of the mice starting from the day of AdV inoculation in both percent weight loss and grams are presented in supplementary figure S2. Collectively, these data suggested that the AdV-hACE2 vector produced morbidity specifically in obese mice. However, we proceeded to test other parameters to determine if differences could be found between lean and obese mice in terms of viral replication, immune response, or lung pathology characteristic of SARS-CoV-2 infection.

**Fig.2:**
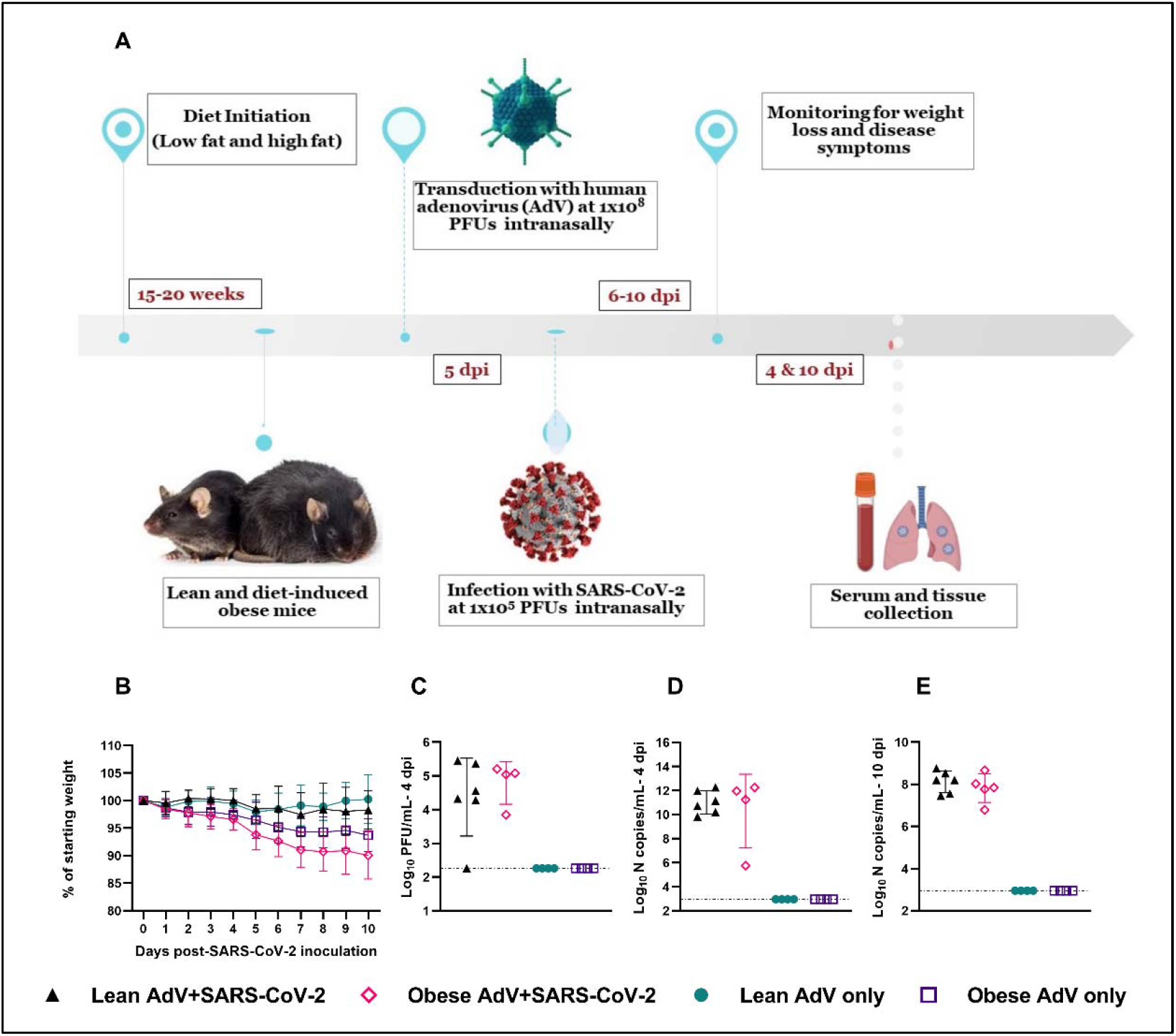
Timeline of the study and disease progression in lean and obese mice post-SARS-CoV-2 infection. **A)** 18-20-week-old male and female C57BL/6N mice received 1×10^8^ PFUs of hACE2-AdV intranasally (day-5) and then 1×10^5^ PFUs SARS-CoV-2 or viral diluent intranasally (day 0). A subset of infected and control mice was euthanized at day 4 and the remaining at day 10 post-SARS-CoV-2 infection and blood and organs were collected for viral titers and histopathology. **B)** Weights were measured daily after SARS-CoV-2 infection and statistical analysis was done on the percent weight change compared to the weight on the day of infection, using 2way-ANOVA with Dunnett’s multiple comparisons test. No significant difference was observed between only adenovirus (AdV) transduced groups and their AdV+SARS-CoV-2 infected counterparts. **C)** Infectious viral titer in the lungs was determined at four dpi. Statistical analysis was done on log transformed data, using Ordinary One-way ANOVA. No significant difference in lung viral titers was observed between lean and obese infected groups (p-value=0.7377). The assay Limit of Detection (L.O.D) was set at 2.255 Log_10_PFUs/mL. **D**) Viral N-copies/mL from RNA extracted from lungs of mice at four dpi, was measured by RT-qPCR. **E)** Viral N-copies/mL from RNA extracted from lungs of mice 10 dpi measured by RT-qPCR. Statistical analysis was done on log transformed data using Ordinary One-way ANOVA. No significant difference in “N” gene copies was observed between lean and obese infected groups at four dpi (p-value=0.8241) or 10 dpi (p-value=0.6052). N copies and L.O.D. were calculated using the standard curve generated from “N-gene RNA standard”. L.O.D. was at a Cq value of 38 or 2.947 Log_10_ N copies/mL.

To measure viral replication, a subset of the mice was euthanized on days 4 and 10 post-SARS-CoV-2 inoculation and infectious virus was quantified in the lung homogenates. As expected, mice receiving only AdV-hACE2 tested negative for SARS-CoV-2 (Fig. 2C). Infectious lung viral titers between lean and obese infected groups were similar (p-value=0.7377) at 4 days post-SARS-CoV-2 infection, and 1 lean and 2 obese mice were negative for infectious virus. We next used RT-qPCR against the N-gene as a more sensitive means to detect viral RNA to determine if the mice became infected with SARS-CoV-2. At 4 days post-SARS-CoV-2 inoculation, all lean mice were positive for the N-gene, while both infectious-virus-negative obese mice were negative, indicating a lack of established infection in these obese mice. Accordingly, we have removed the 2 negative obese mice from Figure 2B-D because they were likely not infected. In Figure 2E, we present RT-qPCR data for the N-gene for lungs collected at 10 days post-SARS-CoV-2 infection; all 6 lean mice, but only 5 obese mice, tested positive for SARS-CoV-2 N-gene by RT-qPCR, and all the mice were negative for infectious virus tested by plaque assays. We present all the data, including the negatives, in supplementary figure S3. No differences were observed between lean and obese groups in N-gene copies/mL at either timepoint. This finding is consistent with previous studies that found no significant difference in infectious viral titers at peak viral replication between lean and obese mice infected with influenza virus [20], several alphaviruses [43] or dengue virus [44].

Severe COVID-19 disease is characterized by low levels of type I interferons (IFN-α and β) and over-production of inflammatory cytokines like interleukin 6 (IL-6) and tumor necrosis factor alpha (TNF-α) [45,46,47,48,49]. Moreover, invasive mechanical ventilation (IMV) was associated with severe obesity in patients infected with SARS-CoV-2, irrespective of age, sex, diabetes, and hypertension [39]. Experiments using obese mice to study the pathogenesis of influenza A virus, another important respiratory virus, showed reduced expression of type I interferons (IFNs), delayed expression of pro-inflammatory cytokines and chemokines [20], severe lung pathology, and increased viral spread, leading to increased mortality [16]. We thus sought to determine the contribution of obesity to cytokine expression and lung pathology following SARS-CoV-2 infection. For cytokine testing, RNA was extracted from lungs collected 4 days post-SARS-CoV-2 infection, and the expression of cytokines IL-6 and IFN-β was measured by RT-qPCR. The relative expression of these cytokines was calculated by the ∆∆Ct method, with hACE-AdV infected lean groups as control and mouse GAPDH as the reference gene. No significant difference in the relative fold-expression of the cytokines above was observed between hACE2-AdV+SARS-CoV-2 infected lean and obese groups at either 4 or 10 dpi (Fig. 3A).

**Fig.3:**
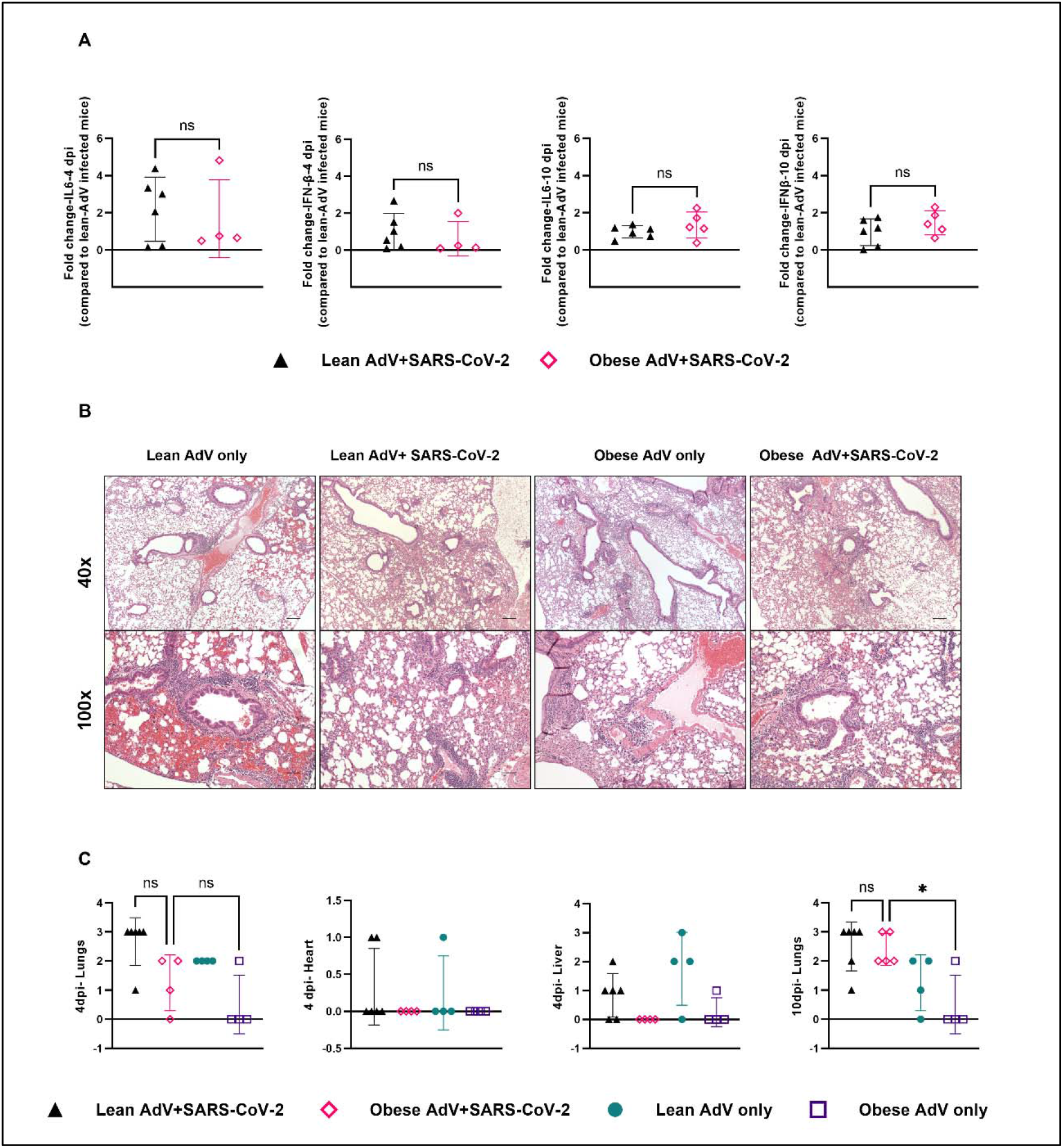
Cytokine mRNA expression and tissue histopathology. **A)** Fold change in gene expression of cytokines, IL-6 and IFN-β at four- and 10-days post-SARS-CoV-2 infection, determined by RT-qPCR, compared to mock-infected mice, and normalized to GAPDH. Statistical analysis was performed by Mann-Whitney test and no significant difference in gene expression was observed between lean and obese groups. **B)** Histopathology slides of the lungs of mice transduced only with AdV-hACE2 and those infected with AdV+SARS-CoV-2 at D4 post-SARS-CoV-2 infection. Images at the top row show a lower magnification (40x) and the bottom row shows medium magnification (100x). No gross pathological differences were observed in lung histopathology between different groups. **C)** Histopathology scores of lungs, heart, and liver four-days post-SARS-CoV-2 infection and for lungs at 10 dpi also. Statistical analysis was done using Ordinary one-way ANOVA with Tukey’s multiple comparisons test. No difference in lung pathology was observed in lean groups infected with AdV+SARS-CoV-2 compared to their obese counterparts at four or 10 dpi (p-value=0.0695 and 0.9971 respectively). The difference between obese mice transduced only with AdV-hACE2 and those inoculated with AdV+SARS-CoV-2 was non-significant at four dpi (p-value=0.5679) but became significant at 10 dpi (p-value=0.0187). The error bars denote SD. No significant difference was observed between the groups in terms of histopathology scores of heart and liver.

We also examined lungs, liver, and heart for histopathological lesions. Lungs were evaluated and scored based on evidence of interstitial inflammation, intra-alveolar hemorrhage, and peribronchiolar and perivascular lymphoid hyperplasia. There was no evidence of interstitial or alveolar septal necrosis or hyaline membrane formation. Livers were evaluated for evidence of lipidosis and single cell necrosis. The hearts were evaluated for evidence of myocardial necrosis and inflammation. Representative lung images collected at 4 dpi are presented in Fig. 3B. The lung histopathology scores were similar for obese mice inoculated with AdV+SARS-CoV-2 compared to their AdV-only transduced counterparts at 4 dpi but became significantly higher for the SARS-CoV-2 infected mice at 10 dpi (Fig. 3C). We could hypothesize that this difference at the later timepoint was due to additional replication of SARS-CoV-2 after 4 dpi and the AdV subsequently being cleared. No differences in the histopathology scores for heart and liver were observed between the groups. Taken together, no major differences were observed between the two groups; however, the weight loss data suggest that the AdV-hACE2 transduction caused an obesity-specific morbidity, thus confounding the study.

The mechanisms underlying the morbidity observed in AdV-hACE2 transduced obese mice are currently unclear. Future work using a control AdV will enable parsing out of vector-related mechanisms vs. ectopic expression of hACE2 will be informative. Replication-defective adenoviruses have been used in many gene therapy applications [50,51] and are known to activate both innate and adaptive immune responses in humans [52,53,54,55]. Further, they have been shown to cause hepatotoxicity in mice due to induction of TNF-α in the liver [56,57] and dose-dependent morbidity in non-human primates [58]. Accordingly, we posit that obese hosts may be more permissive to disease following transduction, possibly due to an overzealous immune response to the vector. The data presented here suggest caution should be taken when using adenovirus vectors in broader populations. Notably, the race for developing a vaccine against SARS-CoV-2 at “pandemic speed” [59] has renewed the interest of researchers in using adenovirus vectors, which makes our findings worth considering.

### Limitations of the study

The major limitation of our study was the use of the adenovirus vector for transduction of mice with the human ACE2 receptor. The use of adenovirus vector was an additional variable that induced morbidity in the obese host, confounding our ability to assess obesity’s impact on SARS-CoV-2 infection in mice, the main objective of our study. Another limitation of this study was the lack of a true mock-treated group, with no AdV or SARS-CoV-2 infection, in the absence of which we cannot quantify the impact of the adenovirus vector versus hACE2 transgene on the obese host. Moreover, these differences could be specific to C57BL/6N mice and we might observe different results with other mouse strains. Despite these constraints, this study provides useful information to the field relating to SARS-CoV-2 mouse models and advises caution when considering the AdV-hACE2 model for nutrition studies.

## Conclusion

The increased risk of grave illness and fatalities in obese people post-SARS-CoV-2 infection underscores the importance of an effective animal model to study the mechanisms underlying worsened disease outcomes. Here, we generated diet-induced obese (DIO) mice and used a previously established adenoviral-transduction model to render them susceptible to SARS-CoV-2 infection. Our results demonstrate that although adenovirus-transduction sensitized obese and lean mice to SARS-CoV-2 infection in their lungs, it also induced morbidity specifically in obese mice. Therefore, this model was not appropriate for studying the relationship between underlying co-morbidities like obesity or diabetes and severe COVID-19. Future studies would benefit from using another mouse model, for example, hACE2 transgenic mice or a mouse-adapted strain of SARS-CoV-2.

## Abbreviations

COVID-19: Coronavirus disease 2019
DIO: diet-induced obesity
ACE2: Angiotensin-converting enzyme 2
SARS-CoV: severe acute respiratory syndrome coronavirus
SARS-CoV-2: severe acute respiratory syndrome coronavirus-2
MERS: Middle East respiratory syndrome coronavirus
hACE2: human angiotensin-converting enzyme
hACE2-Tg: transgenic mice expressing human ACE2
AdV-hACE2: adenovirus vectors expressing hACE2
DMEM: Dulbecco’s Modified Eagle’s Medium
FBS: fetal bovine serum
NEAA: non-essential amino acids
LFD: low-fat diet
HFD: high-fat diet
BSL-3: biosafety level-3
PFUs: plaque forming units
RPMI: Roswell Park Memorial Institute
ViTALS: Virginia Tech Animal Laboratory Services
RT-qPCR: reverse transcription quantitative polymerase chain reaction
IL-6: interleukin-6
IFN-β: interferon-β
L.O.D.: limit of detection
BMI: body-mass index
CHIKV: chikungunya virus
DENV: dengue virus
IFNs: interferons
TNF-α: tumor necrosis factor-α

## Supplementary Information

**Fig.S1:**
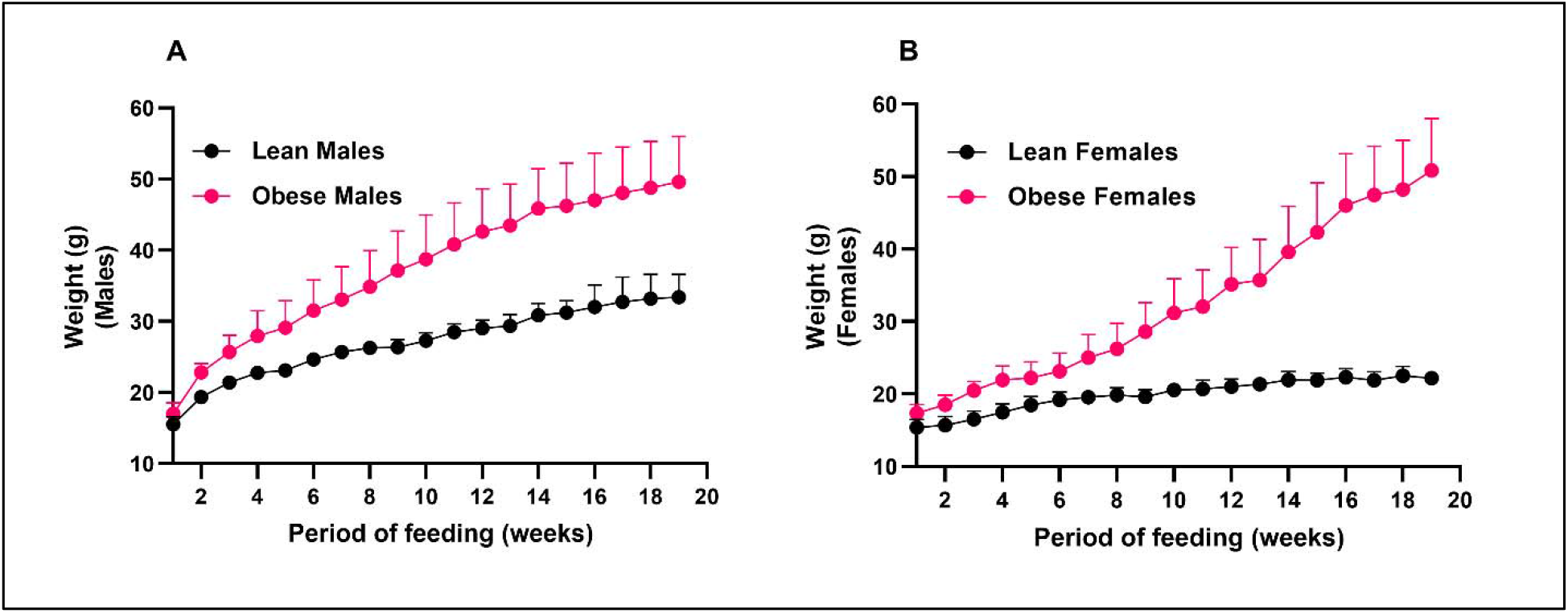
Weight change in males and females on low and high-fat diet. **A)** Weights of C57BL/6N male mice (10/group) fed a high-fat diet (60% fat) or a low-fat diet (10% fat) for 15-20 weeks. Weights were measured weekly after initiating the diets. Statistical analysis was done using 2way-ANOVA with Sidak’s multiple comparisons test. Males on a HFD started showing significant weight gain from four weeks of diet initiation (p-value=0.0114) and gradually became more significant (p-value<0.0001) from seven week onwards. **B)** Weights of C57BL/6N female mice (10/group) fed a high-fat diet (60% fat) or a low-fat diet (10% fat) for 15-20 weeks. Weights were measured weekly after initiating the diets. Statistical analysis was done using 2way-ANOVA with Sidak’s multiple comparisons test. Females on a HFD started showing significant weight gain from six weeks of diet initiation (p-value=0.0104) and gradually became more significant (p-value<0.0001) from eight week onwards.

**Fig.S2:**
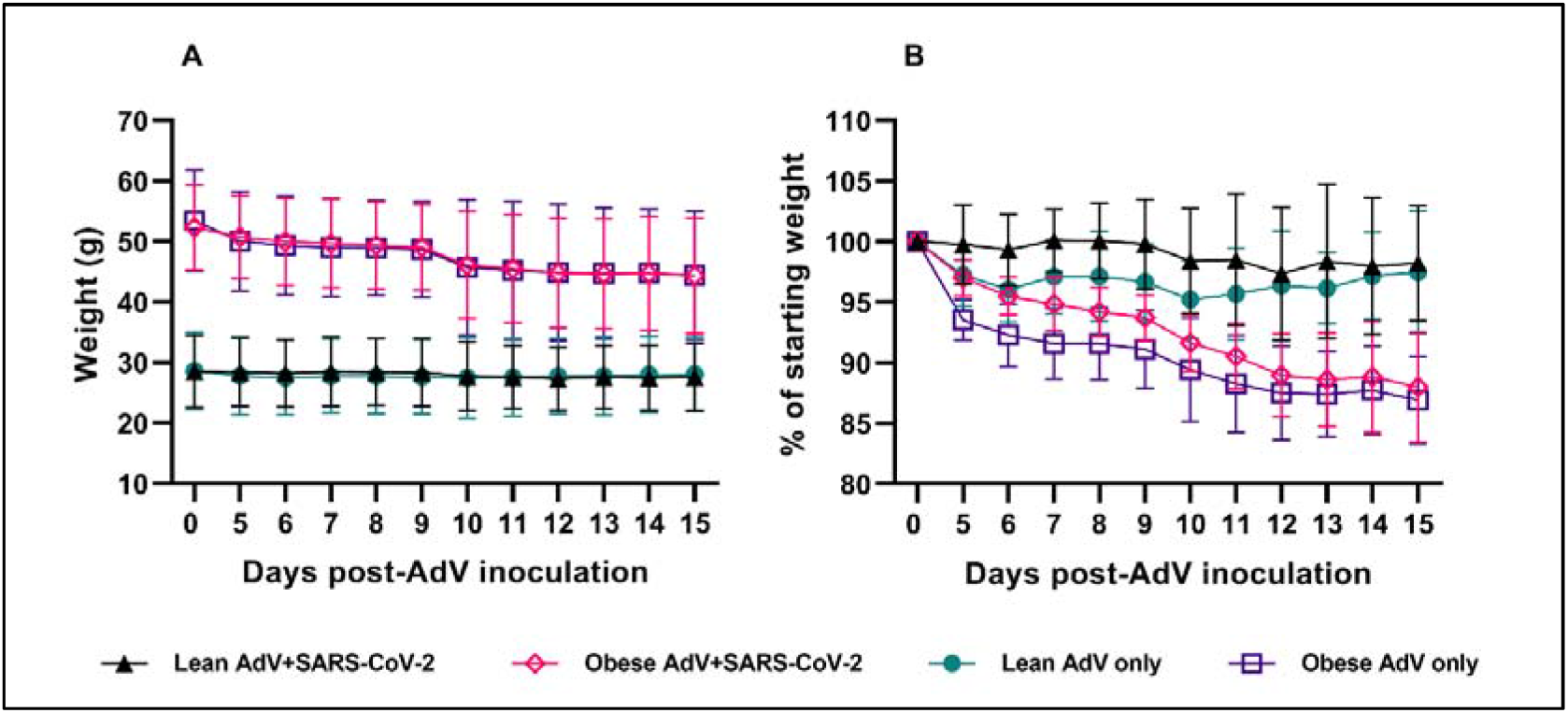
Weight change (actual) and as percent weight loss post-AdV inoculation. **A)**Actual weights of C57BL/6N mice (20/group) post-AdV inoculation (D0), followed by SARS-CoV-2 infection (D5) till the end of experiment (D15). **B)** Weight changes in lean and obese mice inoculated only with AdV vector compared to the ones transduced with the AdV vector and infected with SARS-CoV-2. Statistical analysis was done on the baseline corrected weights, using a 2 way-ANOVA with Dunnett’s multiple comparisons test. A significant decrease in weights was observed in the AdV-only obese mice compared to the lean transduced (only) groups starting at seven dpi (p-value=0.0021) and became highly significant by the end of the experiment (p-value<0.0001).

**Fig.S3:**
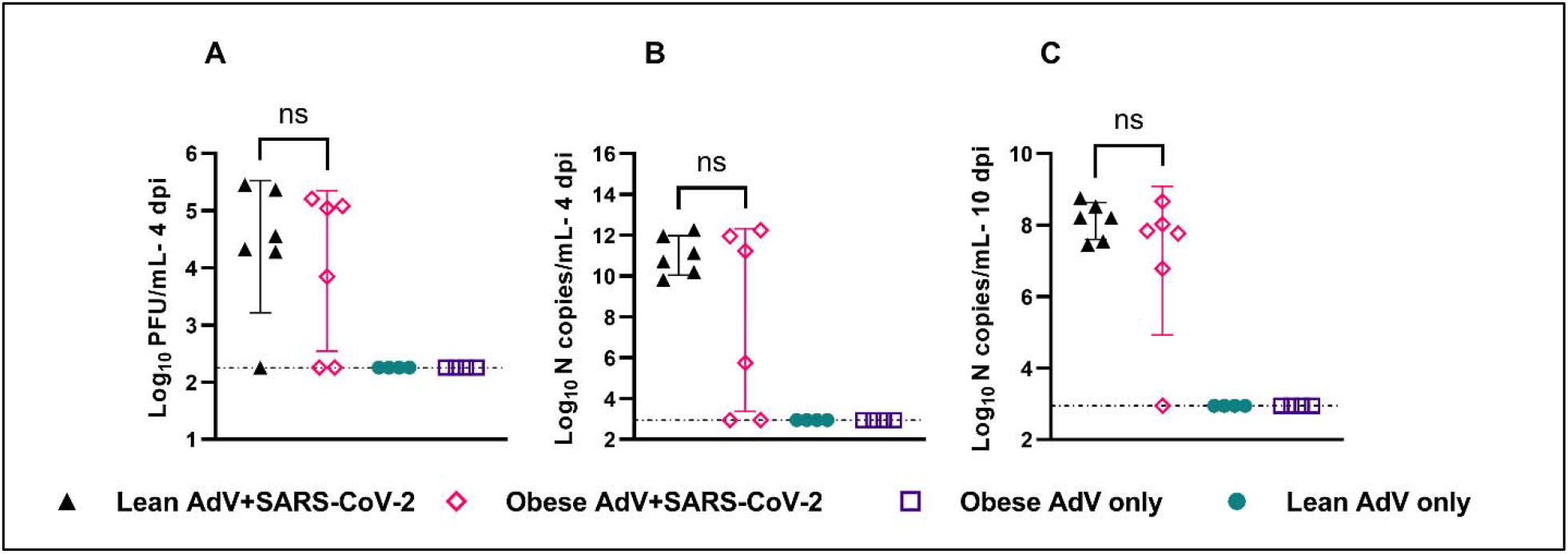
Viral plaque assay titers and N-gene quantification values for all infected mice. **A)** Infectious viral titer in the lungs was determined at four dpi. Statistical analysis was done on log transformed data, using Ordinary One-way ANOVA. No significant difference in lung viral titers was observed between lean and obese infected groups (p-value=0.8224). Samples that tested positive by RT-qPCR, but negative by plaque assays were assigned the L.O.D (2.255 Log_10_PFUs/mL). **B**) Viral N-copies/mL from RNA extracted from lungs of mice at four dpi, was measured by RT-qPCR. **C)** Viral N-copies/mL from RNA extracted from lungs of mice 10 dpi measured by RT-qPCR. Samples with no Cq value were assigned the L.O.D. value (2.947 Log_10_ N copies/mL). Statistical analysis was done on log transformed data using Ordinary One-way ANOVA. No significant difference in “N” gene copies was observed between lean and obese infected groups at four dpi (p-value=0.1198) or 10 dpi (p-value=0.2962). N copies and L.O.D. were calculated using the standard curve generated from “N-gene RNA standard”. L.O.D. was at a Cq value of 38 or 2.947 Log_10_ N copies/mL.

**Table S1:** Caloric information of diets used for the feeding experiments.

| NAME | HIGH FAT DIET (HFD) | LOW FAT DIET (LFD) |
| --- | --- | --- |
| CODE | D12492 | D12450K |
| Protein: | 20 % Kcal | 20 % Kcal |
| Fat: | 60 % Kcal | 10 % Kcal |
| Carbohydrate: | 20 % Kcal | 70 % Kcal |
| Energy density: | 5.21 Kcal/g | 3.82 Kcal/g |

**Table S2:**
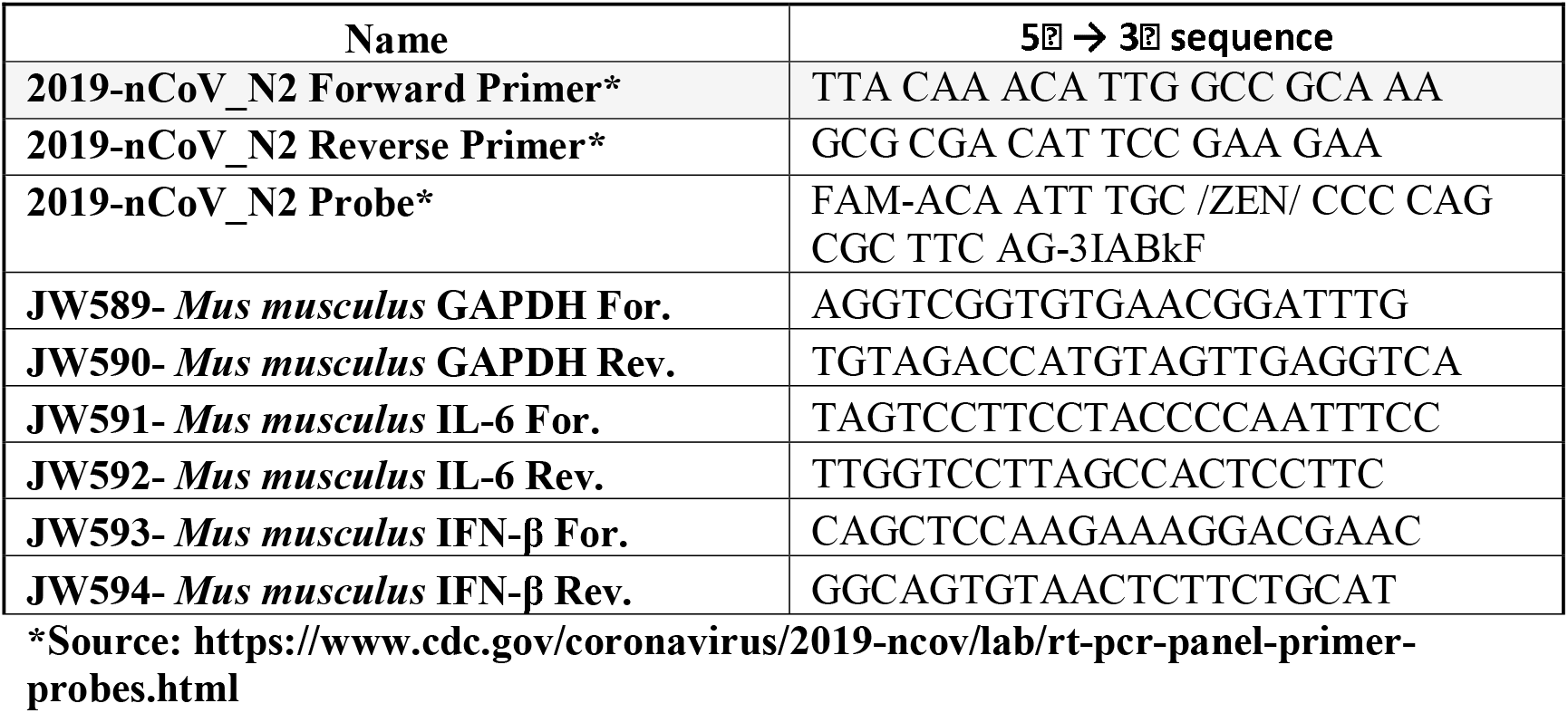
Primers used for RT-qPCR.

## Authors’ contributions

Study concept and design: JWL, PR. Data curation: PR, CC. Formal analysis: PR, JWL. Funding acquisition: JWL. Investigation: PR, CC. Methodology: PR, CC, JWL. Drafting of the manuscript: PR, JWL. Critical revision of the manuscript: all the authors. Statistical analysis: PR, JWL. Histopathology analysis: TLR. All authors have read and approved the final manuscript.

## Availability of data and materials

All data generated or analyzed during this study are included in this published article [and its supplementary information files].

## Ethics approval and consent to participate

All animal handling protocols were approved by the Institutional Animal Care and Use Committee (Protocol #20-060) at Virginia Tech. Consent to participate “not applicable” for this study.

## Consent for publication

Not applicable.

## References

1. Lee N, Hui D, Wu A, Chan P, Cameron P, Joynt GM, et al. A major outbreak of severe acute respiratory syndrome in Hong Kong. N Engl J Med. 2003;348(20):1986–94.

2. Peiris JS, Lai ST, Poon LL, Guan Y, Yam LY, Lim W, et al. Coronavirus as a possible cause of severe acute respiratory syndrome. Lancet. 2003;361(9366):1319–25.

3. Jia HP, Look DC, Shi L, Hickey M, Pewe L, Netland J, et al. ACE2 receptor expression and severe acute respiratory syndrome coronavirus infection depend on differentiation of human airway epithelia. J Virol. 2005;79(23):14614–21.

4. de Wit E, van Doremalen N, Falzarano D, Munster VJ. SARS and MERS: recent insights into emerging coronaviruses. Nat Rev Microbiol. 2016;14(8):523–34.

5. Alagaili AN, Briese T, Mishra N, Kapoor V, Sameroff SC, de Wit E, et al. Middle East Respiratory Syndrome Coronavirus Infection in Dromedary Camels in Saudi Arabia. mBio. 2014;5(2):e00884–14.

6. Memish ZA, Mishra N, Olival KJ, Fagbo SF, Kapoor V, Epstein JH, et al. Middle East respiratory syndrome coronavirus in bats, Saudi Arabia. Emerg Infect Dis. 2013;19(11):1819–23.

7. Zaki AM, van Boheemen S, Bestebroer TM, Osterhaus AD, Fouchier RA. Isolation of a novel coronavirus from a man with pneumonia in Saudi Arabia. N Engl J Med. 2012;367(19):1814–20.

8. WHO. Coronavirus Disease (COVID-19) Dashboard. 2021.

9. Guan W-j, Ni Z-y, Hu Y, Liang W-h, Ou C-q, He J-x, et al. Clinical Characteristics of Coronavirus Disease 2019 in China. New England Journal of Medicine. 2020;382(18):1708–20.

10. Zhou P, Yang X-L, Wang X-G, Hu B, Zhang L, Zhang W, et al. A pneumonia outbreak associated with a new coronavirus of probable bat origin. Nature. 2020;579(7798):270–3.

11. Haimovich AD, Ravindra NG, Stoytchev S, Young HP, Wilson FP, van Dijk D, et al. Development and Validation of the Quick COVID-19 Severity Index: A Prognostic Tool for Early Clinical Decompensation. Ann Emerg Med. 2020;76(4):442–53.

12. de Lusignan S, Dorward J, Correa A, Jones N, Akinyemi O, Amirthalingam G, et al. Risk factors for SARS-CoV-2 among patients in the Oxford Royal College of General Practitioners Research and Surveillance Centre primary care network: a cross-sectional study. The Lancet Infectious Diseases. 2020;20(9):1034–42.

13. Petrilli CM, Jones SA, Yang J, Rajagopalan H, O’Donnell L, Chernyak Y, et al. Factors associated with hospitalization and critical illness among 4,103 patients with Covid-19 disease in New York City. medRxiv. 2020:2020.04.08.20057794.

14. Hui DS, Azhar EI, Kim YJ, Memish ZA, Oh MD, Zumla A. Middle East respiratory syndrome coronavirus: risk factors and determinants of primary, household, and nosocomial transmission. Lancet Infect Dis. 2018;18(8):e217–e27.

15. Van Kerkhove MD, Vandemaele KA, Shinde V, Jaramillo-Gutierrez G, Koukounari A, Donnelly CA, et al. Risk factors for severe outcomes following 2009 influenza A (H1N1) infection: a global pooled analysis. PLoS Med. 2011;8(7):e1001053.

16. O’Brien KB, Vogel P, Duan S, Govorkova EA, Webby RJ, McCullers JA, et al. Impaired wound healing predisposes obese mice to severe influenza virus infection. J Infect Dis. 2012;205(2):252–61.

17. Cocoros NM, Lash TL, DeMaria A, Jr., Klompas M. Obesity as a risk factor for severe influenza-like illness. Influenza Other Respir Viruses. 2014;8(1):25–32.

18. NCHS. Estimates of Diabetes and Its Burden in the United States. 2020. 2020.

19. Engin A. The Definition and Prevalence of Obesity and Metabolic Syndrome. Adv Exp Med Biol. 2017;960:1–17.

20. Smith AG, Sheridan PA, Harp JB, Beck MA. Diet-induced obese mice have increased mortality and altered immune responses when infected with influenza virus. J Nutr. 2007;137(5):1236–43.

21. Honce R, Karlsson EA, Wohlgemuth N, Estrada LD, Meliopoulos VA, Yao J, et al. Obesity-Related Microenvironment Promotes Emergence of Virulent Influenza Virus Strains. mBio. 2020;11(2).

22. Letko M, Marzi A, Munster V. Functional assessment of cell entry and receptor usage for SARS-CoV-2 and other lineage B betacoronaviruses. Nature Microbiology. 2020;5(4):562–9.

23. Wan Y, Shang J, Graham R, Baric RS, Li F. Receptor Recognition by the Novel Coronavirus from Wuhan: an Analysis Based on Decade-Long Structural Studies of SARS Coronavirus. J Virol. 2020;94(7).

24. McCray PB, Pewe L, Wohlford-Lenane C, Hickey M, Manzel L, Shi L, et al. Lethal Infection of K18-*hACE2* Mice Infected with Severe Acute Respiratory Syndrome Coronavirus. Journal of Virology. 2007;81(2):813–21.

25. Yang XH, Deng W, Tong Z, Liu YX, Zhang LF, Zhu H, et al. Mice transgenic for human angiotensin-converting enzyme 2 provide a model for SARS coronavirus infection. Comp Med. 2007;57(5):450–9.

26. Yoshikawa N, Yoshikawa T, Hill T, Huang C, Watts DM, Makino S, et al. Differential Virological and Immunological Outcome of Severe Acute Respiratory Syndrome Coronavirus Infection in Susceptible and Resistant Transgenic Mice Expressing Human Angiotensin-Converting Enzyme 2. Journal of Virology. 2009;83(11):5451–65.

27. Bao L, Deng W, Huang B, Gao H, Liu J, Ren L, et al. The Pathogenicity of SARS-CoV-2 in hACE2 Transgenic Mice. bioRxiv. 2020:2020.02.07.939389.

28. Dinnon KH, Leist SR, Schäfer A, Edwards CE, Martinez DR, Montgomery SA, et al. A mouse-adapted SARS-CoV-2 model for the evaluation of COVID-19 medical countermeasures. bioRxiv. 2020:2020.05.06.081497.

29. Leist SR, Dinnon KH, 3rd, Schäfer A, Tse LV, Okuda K, Hou YJ, et al. A Mouse-Adapted SARS-CoV-2 Induces Acute Lung Injury and Mortality in Standard Laboratory Mice. Cell. 2020;183(4):1070–85.e12.

30. Hassan AO, Case JB, Winkler ES, Thackray LB, Kafai NM, Bailey AL, et al. A SARS-CoV-2 Infection Model in Mice Demonstrates Protection by Neutralizing Antibodies. Cell. 2020;182(3):744–53.e4.

31. Sun J, Zhuang Z, Zheng J, Li K, Wong RL, Liu D, et al. Generation of a Broadly Useful Model for COVID-19 Pathogenesis, Vaccination, and Treatment. Cell. 2020;182(3):734–43.e5.

32. Sia SF, Yan LM, Chin AWH, Fung K, Choy KT, Wong AYL, et al. Pathogenesis and transmission of SARS-CoV-2 in golden hamsters. Nature. 2020;583(7818):834–8.

33. Lee AC-Y, Zhang AJ, Chan JF-W, Li C, Fan Z, Liu F, et al. Oral SARS-CoV-2 Inoculation Establishes Subclinical Respiratory Infection with Virus Shedding in Golden Syrian Hamsters. Cell Reports Medicine. 2020;1(7):100121.

34. Kim YI, Kim SG, Kim SM, Kim EH, Park SJ, Yu KM, et al. Infection and Rapid Transmission of SARS-CoV-2 in Ferrets. Cell Host Microbe. 2020;27(5):704–9.e2.

35. Kutter JS, de Meulder D, Bestebroer TM, Lexmond P, Mulders A, Fouchier RA, et al. SARS-CoV and SARS-CoV-2 are transmitted through the air between ferrets over more than one meter distance. bioRxiv. 2020:2020.10.19.345363.

36. Calhoun PJ, Phan AV, Taylor JD, James CC, Padget RL, Zeitz MJ, et al. Adenovirus targets transcriptional and posttranslational mechanisms to limit gap junction function. Faseb j. 2020;34(7):9694–712.

37. Rogers TF, Zhao F, Huang D, Beutler N, Burns A, He WT, et al. Isolation of potent SARS-CoV-2 neutralizing antibodies and protection from disease in a small animal model. Science. 2020;369(6506):956–63.

38. Livak KJ, Schmittgen TD. Analysis of relative gene expression data using real-time quantitative PCR and the 2(-Delta Delta C(T)) Method. Methods. 2001;25(4):402–8.

39. Simonnet A, Chetboun M, Poissy J, Raverdy V, Noulette J, Duhamel A, et al. High Prevalence of Obesity in Severe Acute Respiratory Syndrome Coronavirus-2 (SARS-CoV-2) Requiring Invasive Mechanical Ventilation. Obesity (Silver Spring). 2020;28(7):1195–9.

40. Bassetti M, Parisini A, Calzi A, Pallavicini FM, Cassola G, Artioli S, et al. Risk factors for severe complications of the novel influenza A (H1N1): analysis of patients hospitalized in Italy. Clin Microbiol Infect. 2011;17(2):247–50.

41. Balanzat AM, Hertlein C, Apezteguia C, Bonvehi P, Cámera L, Gentile A, et al. An analysis of 332 fatalities infected with pandemic 2009 influenza A (H1N1) in Argentina. PLoS One. 2012;7(4):e33670.

42. Huttunen R, Syrjänen J. Obesity and the risk and outcome of infection. Int J Obes (Lond). 2013;37(3):333–40.

43. Weger-Lucarelli J, Carrau L, Levi LI, Rezelj V, Vallet T, Blanc H, et al. Host nutritional status affects alphavirus virulence, transmission, and evolution. PLOS Pathogens. 2019;15(11):e1008089.

44. Chuong C, Bates TA, Akter S, Werre SR, LeRoith T, Weger-Lucarelli J. Nutritional status impacts dengue virus infection in mice. BMC Biology. 2020;18(1):106.

45. Hadjadj J, Yatim N, Barnabei L, Corneau A, Boussier J, Smith N, et al. Impaired type I interferon activity and inflammatory responses in severe COVID-19 patients. Science. 2020;369(6504):718–24.

46. Huang C, Wang Y, Li X, Ren L, Zhao J, Hu Y, et al. Clinical features of patients infected with 2019 novel coronavirus in Wuhan, China. The Lancet. 2020;395(10223):497–506.

47. Mehta P, McAuley DF, Brown M, Sanchez E, Tattersall RS, Manson JJ. COVID-19: consider cytokine storm syndromes and immunosuppression. The Lancet. 2020;395(10229):1033–4.

48. Wen W, Su W, Tang H, Le W, Zhang X, Zheng Y, et al. Immune cell profiling of COVID-19 patients in the recovery stageby single-cell sequencing. Cell Discovery. 2020;6(1):31.

49. Chua RL, Lukassen S, Trump S, Hennig BP, Wendisch D, Pott F, et al. COVID-19 severity correlates with airway epithelium–immune cell interactions identified by single-cell analysis. Nature Biotechnology. 2020;38(8):970–9.

50. Crystal RG. Transfer of genes to humans: early lessons and obstacles to success. Science. 1995;270(5235):404–10.

51. Thacker EE, Nakayama M, Smith BF, Bird RC, Muminova Z, Strong TV, et al. A genetically engineered adenovirus vector targeted to CD40 mediates transduction of canine dendritic cells and promotes antigen-specific immune responses in vivo. Vaccine. 2009;27(50):7116–24.

52. Juillard V, Villefroy P, Godfrin D, Pavirani A, Venet A, Guillet JG. Long-term humoral and cellular immunity induced by a single immunization with replication-defective adenovirus recombinant vector. Eur J Immunol. 1995;25(12):3467–73.

53. Yang Y, Li Q, Ertl HC, Wilson JM. Cellular and humoral immune responses to viral antigens create barriers to lung-directed gene therapy with recombinant adenoviruses. J Virol. 1995;69(4):2004–15.

54. Adesanya MR, Redman RS, Baum BJ, O’Connell BC. Immediate inflammatory responses to adenovirus-mediated gene transfer in rat salivary glands. Hum Gene Ther. 1996;7(9):1085–93.

55. Worgall S, Wolff G, Falck-Pedersen E, Crystal RG. Innate immune mechanisms dominate elimination of adenoviral vectors following in vivo administration. Hum Gene Ther. 1997;8(1):37–44.

56. Engler H, Machemer T, Philopena J, Wen S-F, Quijano E, Ramachandra M, et al. Acute hepatotoxicity of oncolytic adenoviruses in mouse models is associated with expression of wild-type E1a and induction of TNF-α. Virology. 2004;328(1):52–61.

57. Varnavski AN, Calcedo R, Bove M, Gao G, Wilson JM. Evaluation of toxicity from high-dose systemic administration of recombinant adenovirus vector in vector-naive and pre-immunized mice. Gene Ther. 2005;12(5):427–36.

58. Schnell MA, Zhang Y, Tazelaar J, Gao GP, Yu QC, Qian R, et al. Activation of innate immunity in nonhuman primates following intraportal administration of adenoviral vectors. Mol Ther. 2001;3(5 Pt 1):708–22.

59. Lurie N, Saville M, Hatchett R, Halton J. Developing Covid-19 Vaccines at Pandemic Speed. New England Journal of Medicine. 2020;382(21):1969–73.

